# Heteromerization of alkaline-sensitive two-pore domain potassium channels

**DOI:** 10.1101/2021.11.08.467666

**Authors:** Lamyaa Khoubza, Eun-Jin Kim, Franck C. Chatelain, Sylvain Feliciangeli, Dawon Kang, Florian Lesage, Delphine Bichet

## Abstract

Two-pore domain (K_2P_) potassium channels are active as dimers. They produce inhibitory currents regulated by a variety of stimuli. Among them, TALK1, TALK2 and TASK2 form a subfamily of structurally related K_2P_ channels stimulated by extracellular alkalosis. The human genes encoding them are clustered on chromosomal region 6p21. They are expressed in different tissues including the pancreas. By analyzing single cell transcriptomic data, we show that these channels are co-expressed in insulin-secreting pancreatic β cells. By different approaches we show that they form functional heterodimers. Heteromerization of TALK2 with TALK1 or with TASK2 endorses TALK2 with sensitivity to extracellular alkalosis in the physiological range. The association of TASK2 with TALK1 and TALK2 increases their unitary conductance. These results provide a new example of heteromerization in the K_2P_ channel family expanding the range of their potential physiological and pathophysiological roles.

## Introduction

Two-pore domain K^+^ channels (K_2P_ channels) produce inhibitory background potassium currents regulated by a variety of chemical and physical stimuli (1–3). They are active as dimers of subunits, with each subunit containing 4 membrane-spanning segments (M1 to M4) and 2 pore-domains (P1 and P2). Their N- and the C-termini are intracellular. The TWIK1-related alkalinization-activated K^+^ channel 1 (TALK1), TWIK1-related alkalinization-activated K^+^ channel 2 (TALK2) and TWIK1-related acid-sensitive K^+^ channel 2 (TASK2) form a subgroup of structurally and functionally related K_2P_ channels (Fig. 1A). TALK1 shares 44% sequence identity with TALK2 and 39% with TASK2, whereas TALK2 and TASK2 share 37% sequence identity (4,5) (Fig. 1A). They produce K^+^ currents that share the particularity to be stimulated by alkalinization of the extracellular medium (4–6). They are also activated by NO and ROS and intracellular pH (6). Recent research suggests a regulatory function of TALK1 for glucose-dependent β-cell second-phase insulin and δ-cell somatostatin secretion (7,8). The role of TALK1 in glucose-stimulated insulin secretion (GSIS) is further supported by the fact that islet β-cells from TALK1 knockout (KO) mice exhibit increased *V*_m_ depolarization, augmented Ca^2+^ influx, and elevated second phase GSIS (7). In human, the TALK1 polymorphism rs1535500, which results in a gain-of-function (A277E), has been linked to an increased risk of type 2 diabetes (9,10). More recently, another gain-of-function mutation in TALK1 (L114P) was identified in a family with maturity-onset diabetes of the young (MODY). Both mutations resulted in reduced GSIS due to impaired intracellular Ca^2+^ homeostasis (7,11). Very little is known about the physiological function of TALK2, as this channel is absent in mice. A gain-of-function mutation in TALK2 (G88R) was described in a human patient with another mutation in a sodium channel gene (*SCNA5*) with a cardiac phenotype suggesting that TALK2 may act as a modifier of cardiac arrhythmia (12). In contrast to TALK2, TASK2 channel has been implicated in several functions mostly in brain and kidneys. TASK2 is involved in central oxygen chemoreception (13), cell volume regulation and bicarbonate reabsorption in the kidney (14,15). A loss-of-function mutation (T180P) with dominant-negative effect on wild-type TASK2 has been reported with higher frequency among patients predisposed to Balkan Endemic Nephropathy (BEN), a chronic kidney disease (16,17).

**Fig. 1.**
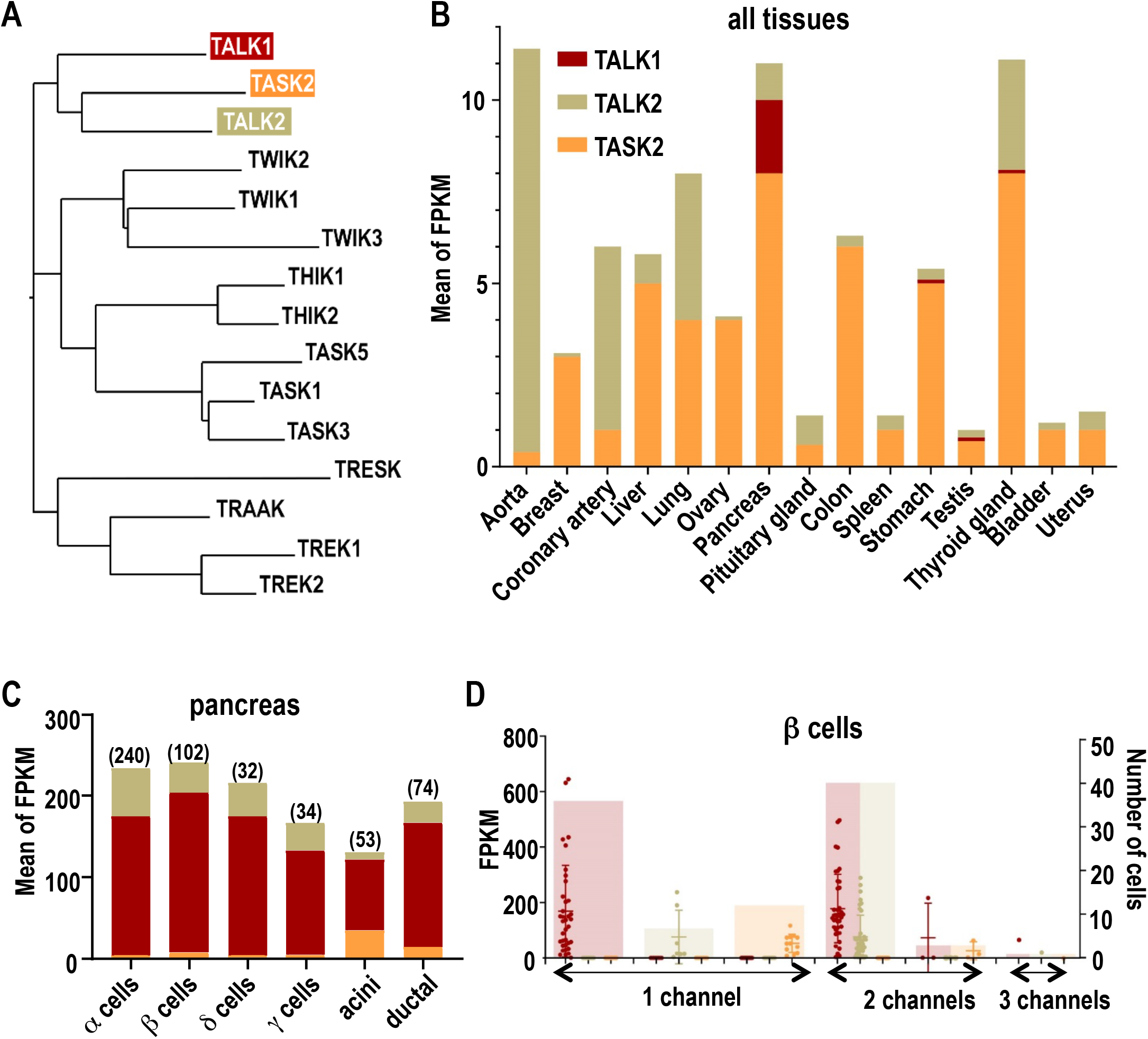
Expression pattern of TALK1, TALK2 and TASK2 channels from RNA-Seq database analysis. (A) TALK1, TALK2 and TASK2 position within the K2P phylogenetic tree of human K2P channels. (B) Human tissue distribution. Expression levels are expressed as the mean of Fragments Per Kilobase of transcript, per Million mapped reads (FPKM). mRNA-seq datasets were extracted from the EMBL database in a study of 1,641 samples across 43 tissues from 175 individuals (GTEx consortium). (C) Expression levels in various human pancreatic cells population including α, β, γ, ε acinar, and ductal cells. Single-cell transcriptomic data were from 2,209 cells. The number of cells analyzed for each population is indicated. (D) Expression in individual human β cells. Cells are divided into 3 groups displaying cells expressing a unique TALK/TASK subunit (1 channel), two subunits (2 channels) and three subunits (3 channels).

Heterodimerization between pore-forming subunits occurs within the K_2P_ channel family (see (18) for recent review). Heterodimeric channels often exhibit unique electrophysiological and pharmacological properties compared to the corresponding homodimers. By mixing and matching subunits, heteromerization increases functional diversity and contributes to pharmacological heterogeneity in the K2P channel family. The first demonstration of heteromerization was between TASK1 and TASK3 (19). Then the following heterodimers were reported: THIK1/THIK2, TREK1/TREK2, TREK1/TRAAK and TREK2/TRAAK (20–23). Heterodimers between members of different subgroups of K_2P_ channels have also been described: TWIK1/TASK1 or 3, TWIK1/TREK1 or 2, TASK1/TALK2, TRESK/TREK1,2 (24–29). Although some of these intergroup heterodimers have been confirmed by independent studies, others remain unclear as they were not detected by other studies (18).

The genes encoding TALK1 (*KCNK16*), TALK2 (*KCNK17*) and TASK2 (*KCNK5*) are located in the same chromosomal region (6p21), *KCNK16* being separated from *KCNK17* by less than 1 kb (4). This genetic clustering suggests that their transcription may be coordinated (at least partially), and that these channel subunits may associate to form heterodimeric channels. TALK1 is expressed in pancreatic cells and gastric somatostatin cells (6,8). Low signal expression was also detected in the small intestine and stomach (30). TALK1, TALK2 and TASK2 are all expressed in the pancreas. Unlike TALK1, TALK2 and TASK2 are expressed in other tissues. TALK2 is present in placenta, lung, liver, small intestine, heart and aorta and, to a lesser extent in the brain (31,32). TASK2 is abundant in the kidney, salivary glands, colon and immune T cells. In the brain, TASK2 expression is restricted to specific regions such as brainstem nuclei, hippocampus and cerebellum (33–36).

Here we reexamine the coexpression of TALK1, TALK2 and TASK2 at the single-cell level and provide evidence of a physical and functional interaction between these subunits. Their heteromerization produces channels with unique pH sensitivity and single-channel properties.

## Results

### TALK1, TALK2 and TASK2 subunits are coexpressed in tissues and cells

In the previous studies reporting the identification and characterization of the TALK1, TALK2 and TASK2 channels, their tissue distribution was analyzed by Northern blots or RT-PCR (4,5,8,30–36). These results cannot be used for comparing their relative expression levels in the tissues in which they are expressed, as the experimental conditions (blots and probes for Northern blot analysis, and cDNA templates and number of cycles for PCR) were not the same. We reexamined the distribution of these channels by taking advantage of data collected using transcriptomics that allow such a comparison, in tissues but also in cells.

We first analyzed the expression of TALK1, TALK2 and TASK2 in the EMBL expression atlas database (https://www.ebi.ac.uk/gxa/). The analysis of an RNA-seq dataset of 53 human tissues from the Genotype-Tissue Expression (GTEx) project shows that TASK2 and TALK2 are present in most tissues while TALK1 has a more restricted distribution in pancreas, stomach, testis and thyroid gland (Fig. 1B). Some tissues preferentially express one channel, such as TALK2 in aorta and coronary artery or TASK2 in the liver, ovary, colon and bladder. In the lung, TALK2 and TASK2 are expressed at the same level. Only the pancreas expresses significant levels of the three channels (Fig. 1B).

Because tissues are composed of different cell types that have the potential to express these channels at different levels, we next analyzed their expression in the major cell types of the pancreas. Data were extracted from a previous RNA-seq study that profiled human pancreatic cells using single-cell transcriptomics (37). We analyzed channel expression in endocrine (α, β, γ, δ, ε) and exocrine (acinar and ductal) cells (Fig. 1C). All cell types express TALK1, TALK2 and TASK2. However, TASK2 was found to be less expressed, relative to TALK1 and TALK2 expression, than expected from the EMBL expression atlas database (Fig. 1B). This may be due to the expression of TASK2 in another pancreatic cell type or to inter individual variations between the samples analyzed in these two studies (variations related to age, gender or ethnicity).

Because RNA-seq was performed at the single cell level (37), we were then able to analyze channel expression in individual cells of a given cell type. Figure 1D shows the analysis carried out for insulin-secreting β cells. TALK1 was detected in 80 out of the 99 cells studied (Fig. 1D). TALK2 was detected in 48 cells and TASK2 in 16 cells. Interestingly, both TALK1 and TALK2 were detected in 40 cells, and TALK1 and TASK2 in 3 cells. TALK1, TALK2 and TASK2 were co-detected in 1 cell out of 99 analyzed.

Taken together these results show that TALK1, TALK2 and TASK2 have overlapping tissues distribution, and that they can be co-expressed at the cell level in the pancreatic β cells. This suggested, by inference with what we have previously shown for the TREK and THIK subfamilies, that TALK1, TALK2 and TASK2 could associate to form heterodimers.

### TALK subunits physically interact in mammalian cells

A physical interaction between TALK1, TALK2 and TASK2 was first tested using *in situ* proximity ligation assay (PLA) in cultured mammalian cells (Fig. 2A). This technique uses primary antibodies that bind to tags present on channel subunits. Primary antibodies are labeled with secondary antibodies containing complementary DNA strands. When antibodies are in close proximity (30-40 nm apart), the two strands hybridize, enabling subsequent DNA amplification by PCR. This amplification product is detected as a red fluorescent signal. Each pair of primary antibodies was tested for specificity (Fig. 2B). TALK1-FLAG subunit, cloned into the pIRES-EGFP vector for green fluorescence coexpression, was expressed with TALK1-HA or TALK1-V5 for positive controls (formation of homodimers) and with the distantly related K^+^ channel subunits K_V_1.2-HA or K_IR_2.1-V5 for negative controls. The percentage of positive cells was calculated by determining the ratio of cells with a positive signal (red) among the hundreds of transfected cells (green), from 3 to 7 independent experiments (Fig. 2C).

**Fig. 2.**
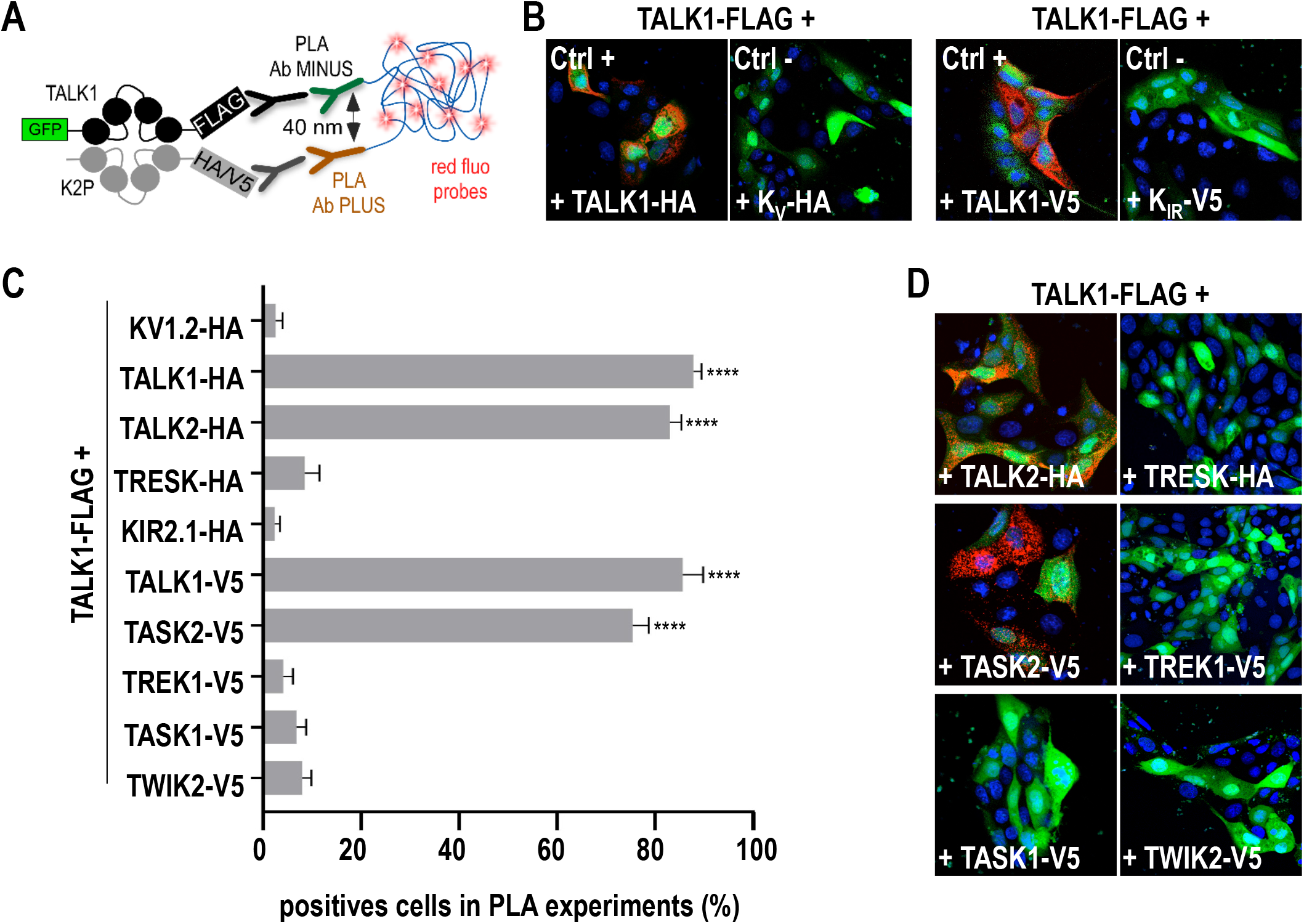
Physical interaction of TALK1, TALK2 and TASK2 by proximity ligation assay (PLA). Tagged K2P channels were expressed in MDCK cells. (A) When the two PLA probes are close enough (<40 nm), a ligation generates circle DNA. This DNA is then amplified by a polymerase, and complementary fluorescent nucleotides are incorporated giving a red positive PLA signal. (B) Positive and negative controls. (C) Quantification of the PLA signal. Percentage of cells is the number of PLA-positive cells (red) relative to the total number of transfected cells (green). The number of cells observed per condition is >100 per condition, and each condition was repeated at least 3 times. Data were analyzed by using unpaired t-test: ****P < 0.0001 versus negative control. (D) Representative microscopy images used for PLA quantification.

As expected, most cells showed a positive PLA signal when the TALK1 homodimer is formed, i.e. between TALK1-FLAG and TALK1-HA (88%) or TALK1-FLAG and TALK1-V5 (85%), whereas less than 3% of cells were positives when TALK1-FLAG was coexpressed with K_V_1.2-HA or KIR2.1-V5, demonstrating the absence of interaction (Fig. 2B, C). The same difference was observed when TALK1-FLAG was coexpressed with TALK2-HA (83%) or TASK2-V5 (75%) compared to TREK1-V5 (4%), TASK1-V5 (7%), TRESK-HA (9%) or TWIK2-V5 (8%) (Fig. 2C, D). These results show that TALK1 can form heterodimers with TALK2 and TASK2 but not with members of other K_2P_ channel subfamilies.

### A dominant negative TALK1 subunit decreases TALK2 and TASK2 currents in Xenopus oocytes

To confirm the interaction between TALK1, TALK2 and TASK2, we used a functional method based on channel poisoning with a non-functional subunit in *Xenopus* oocytes (Fig. 3A). The same dominant negative (DN) strategy was used to demonstrate heteromerization in TREK and THIK K_2P_ channel subfamilies (20,21). In TALK1, we replaced the glycine residue at position 110 with a glutamate residue (G110E). This mutation in the pore domain of the channel leads to a loss of function. As expected, TALK1-G110E produced no measurable current (0.4 μA ± 0.1 *versus* 2.2 μA ± 0.3 for TALK1, at +60 mV, n=9). To test its effect on TALK1, TALK1-G110E and TALK1 cRNAs were injected in equal amount. Representative current traces (Fig. 3C) and normalized mean current amplitudes (Fig. 3B) show that TALK1 is largely inhibited by TALK1-G110E (0.3 μA ± 0.07, n=12) confirming the dominant negative action of TALK1-G110E, renamed TALK1^DN^.

**Fig. 3.**
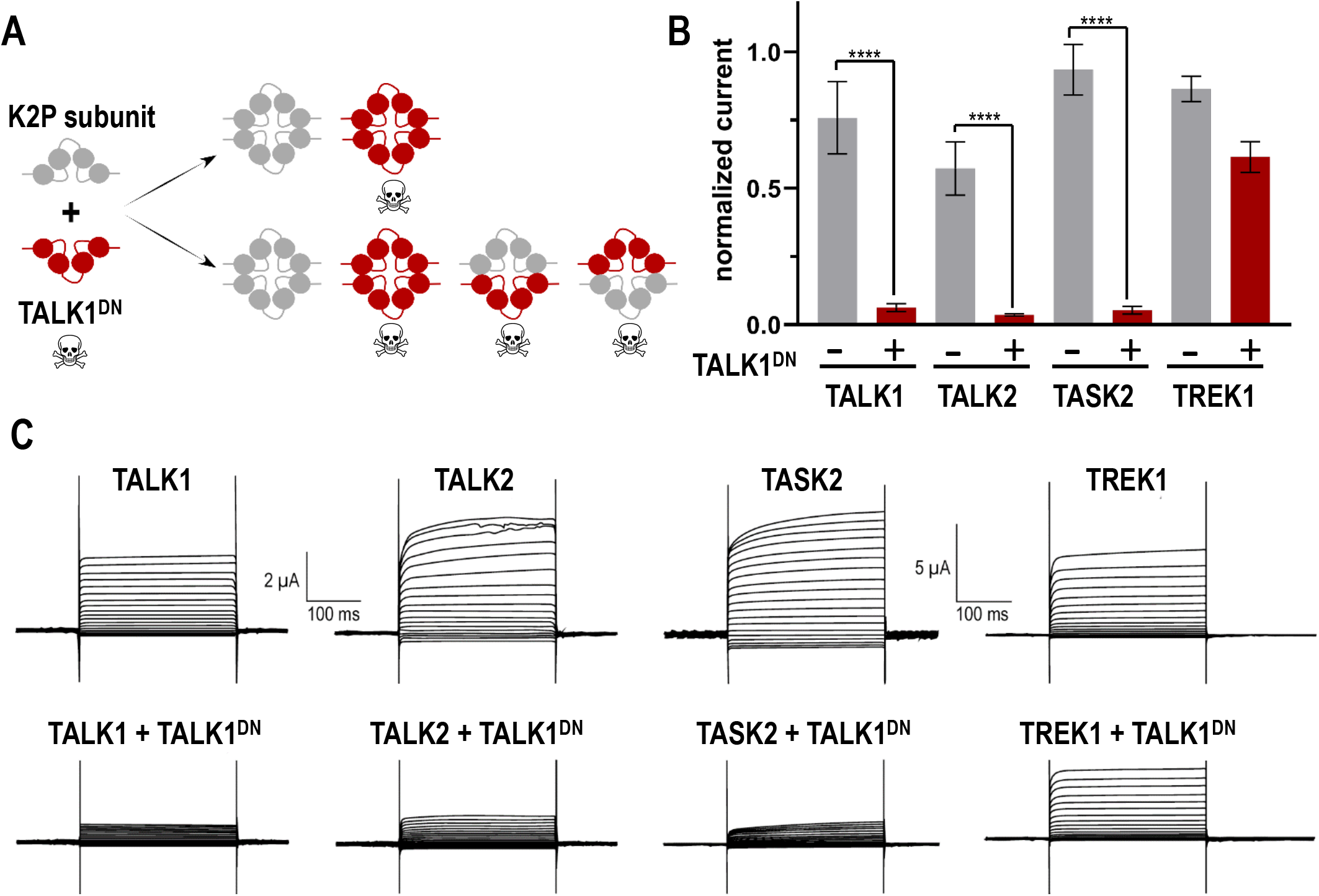
Heterodimerization of TALK1, TALK2 and TASK2 shown by current inhibition by a dominant negative TALK1 (TALK1DN) coexpressed in Xenopus oocytes. (A) Method: a non-functional subunit (red) bearing a mutation in the pore domain (G108E) is coinjected in a 1:1 ratio with a functional K2P subunit. In the absence of heterodimerization, the current is not affected by TALK1DN. If TALK1DN and the coexpressed subunit associate, then the heterodimers are nonfunctional and the current decreases by at least 75%. (B) Normalized steady-state average current amplitude (mean ± S.E.) at 0 mV current for oocytes expressing K2P subunit alone or with TALK1DN, as shown. Data were analyzed by using unpaired t-test: ****P < 0.0001 versus negative control. (C) Representative traces obtained from oocytes expressing TALK1, TALK2, TASK2, and TREK1 in the absence or presence of TALK1DN. Currents were recorded at membrane potentials ranging from −120 mV to +60 mV from a holding potential of −80 mV in 10 mV increments.

We next coexpressed TALK1^DN^ with other K_2P_ subunits. In oocytes, expression of TALK2, TASK2 and TREK1 produced currents as expected (Fig. 3C). At +60 mV, the mean current amplitudes were 3.5 μA ± 0.6 (n=7), 7.2 μA ± 0.7 (n=7) and 7.8 ± 0.5 μA (n=7), respectively. Coexpression of TALK1^DN^ with TALK2 and TASK2 caused a dramatic decrease in current amplitude (Fig. 3C). The decrease reached 87% for TALK2 and 94% for TASK2, values that are comparable to the effect on TALK1 (86%). These values are consistent with the 75% decrease in current amplitude expected for 1:1 ratio of negative dominant to functional subunit (Fig. 3A). The inhibition of TREK1 by TALK1^DN^ is much more limited (30%), suggesting a competitive effect on protein expression rather than heteromerization (Fig. 3B, C). Taken together, the results of this electrophysiological approach confirm the formation of heteromers between TALK1 and TALK2 or TASK2.

### TALK heterodimers exhibits unique pH sensitivity

We then asked whether heteromerization between TALK1 and TALK2 or TASK2 produces heteromeric channels with distinct properties. An important characteristic of these channels is their sensibility to the extracellular pH. They are stimulated by alkalinization in the range of pH 7.5-10 (4,5,38). This sensitivity involves a pH sensor containing a titrable residue located in the extracellular P2-M4 loop: R242 in TALK1, K242 in TALK2 and R224 in TASK2 (39–41) (Fig. 4A). The substitution of this basic residue by a neutral residue confers insensitivity to alkalinization. To compare the pH sensitivity of TALK1, TALK2 and TASK2 under the same experimental conditions, they were expressed in *Xenopus* oocytes. The currents were measured at 0 mV, normalized to the currents measured at pH 10, and plotted as a function of the extracellular pH (Fig. 4B). As previously reported, each homodimeric channel has a unique profile of sensitivity to external pH, that is different from each other and from other pH-sensitive K2P channels including TASK and TREK channels (38). TALK1 is open above pH6 with a biphasic pH-dependence curve (4,41). pH-dependence curve for TASK2 is shifted to more alkaline values from pH6 to 8 but then almost superpose with TALK1 second phase from pH8 to pH10 (Fig. 4B). The estimated pK_1_/_2_ for the effect of pH on TASK2 is near 8, close to the values reported in mammalian cells (5,41,42). The pH-dependence of TALK2 is significantly different from that of TALK1 and TASK2, with a shift to more alkaline values. As previously reported for this channel, almost no current is detected at physiological pH, then the current starts to increase sharply from pH 8 without apparent saturation at pH 10 (4) (Fig. 4B).

**Fig. 4:**
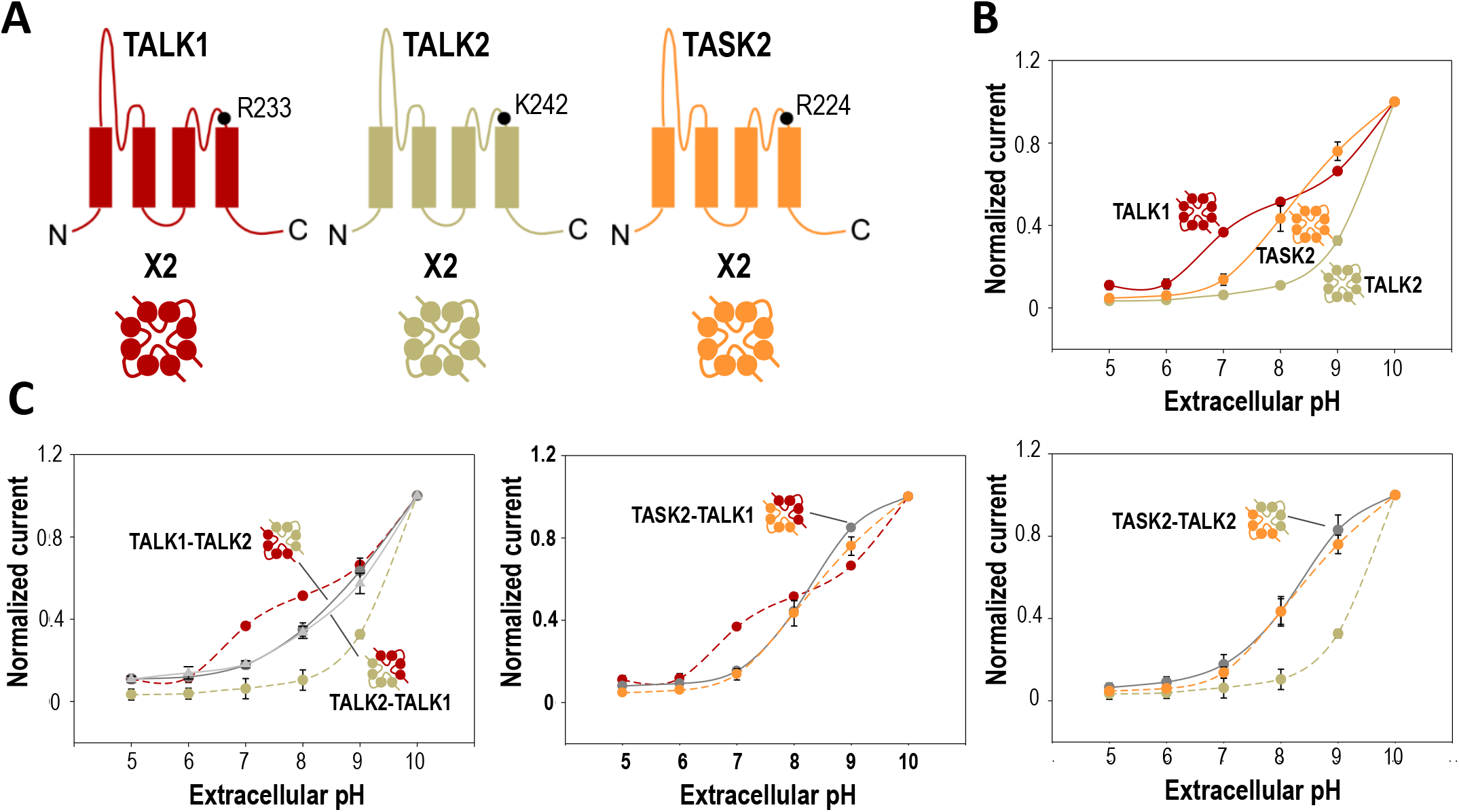
Sensitivity to extracellular pH of homomeric and heteromeric TALK/TASK channels expressed in Xenopus oocytes. (A) Overall topology of TALK/TASK2 subunits. Each subunit comprises 4 transmembrane domains, two extracellular pore regions and cytoplasmic N- and C-ter. Basic residues involved in extracellular pH sensing are shown. (B, C) pH-dependence curves measured with subunits expressed alone (B), or in covalently linked tandem (C). Values represent means ± SEM (n=10) of currents measured at 0 mV, normalized to the currents measured at pH 10, and plotted against extracellular pH. (C) Comparison of the extracellular pH-dependence curves. Dashed lines, pH-dependence curves of homomeric channels as in (B).

To study the pH-sensibility of heterodimeric channels, we designed cDNAs for the expression of tandem chimeras in which two different subunits are covalently linked through the C-ter of the first subunit and the N-ter of the second. This method allows access to currents produced by pure heterodimeric channels rather than by a mixture of homo- and heterodimers, as observed when the two subunits are expressed separately. For TASK1-TASK3 tandems, differences in electrophysiological behavior have been reported depending on the order of the two subunits in the chimeras (43). We therefore designed and expressed cDNAs for both Td-TALK1/TALK2 and Td-TALK2/TALK1 channels. For tandems containing TASK2, only Td-TASK2/TALK1 and Td-TASK2/TALK2 yielded measurable currents, as the tandems with the opposite orientation (Td-TALK1/TASK2 and Td-TALK2/TASK2) produced no current, suggesting that a physical constraint on the N-ter of TASK2 could alter channel assembly and folding, or trafficking to the plasma membrane. The pH-dependence of the functional tandems was compared with the pH-dependence of the corresponding homomers (Fig. 4C). Td-TALK1-TALK2 and Td-TALK2-TALK1 show an intermediate pH-dependence compared to TALK1 and TALK2. The heterodimeric channels are inhibited at acidic pH and activated at basic pH with a dependence curve resembling the monophasic one of TALK2 but shifted to more acidic values. As for TALK2, the current increase is not saturating even under very basic conditions (pH 10). For Td-TASK2/TALK1 and Td-TASK2/TALK2, the pH-dependence curves are very similar and overlap with the curve for TASK2 (Fig. 4C). Compared with TALK2 alone, the combination of TASK2 with TALK2 results in more current around physiological values.

These results show that heteromerization has a very significant effect on the extracellular pH sensitivity of the TALK/TASK2 channels: association of TALK1 with TALK2, or of one of these subunits with TASK2 produces heterodimeric channels that behave like TASK2.

### Single channel properties of the TALK tandems

We next compared the single channel properties of homodimers and heterodimers in mammalian cells. Cell-attached patch recordings from HEK293 cells expressing TALK1, TALK2 and TASK2 show typical single-channel openings (Fig. 5). Under symmetric K^+^ conditions (150 mM KCl, pH_o_ 7.4), TALK1 is active over the entire range of membrane potentials and shows very brief openings. The mean open time was less than 0.3 ms (0.2 ± 0.1 ms), close to reported values (Han et al., 2003; Kang & Kim, 2004) (Fig. 5A). The unitary conductance of TALK1 is 22 ± 1 pS at −80 mV and 9.8 ± 1 pS at +80 mV. The mean open time of TALK2 is 0.8 ± 0.1 ms (−80 mV), a value significantly higher than that of TALK1. The unitary conductance of TALK2 is 41 ± 3 pS at −80 mV and 14 ± 2 pS at +80 mV, values that are also significantly higher than those of TALK1 (Fig. 5A).

**Fig. 5:**
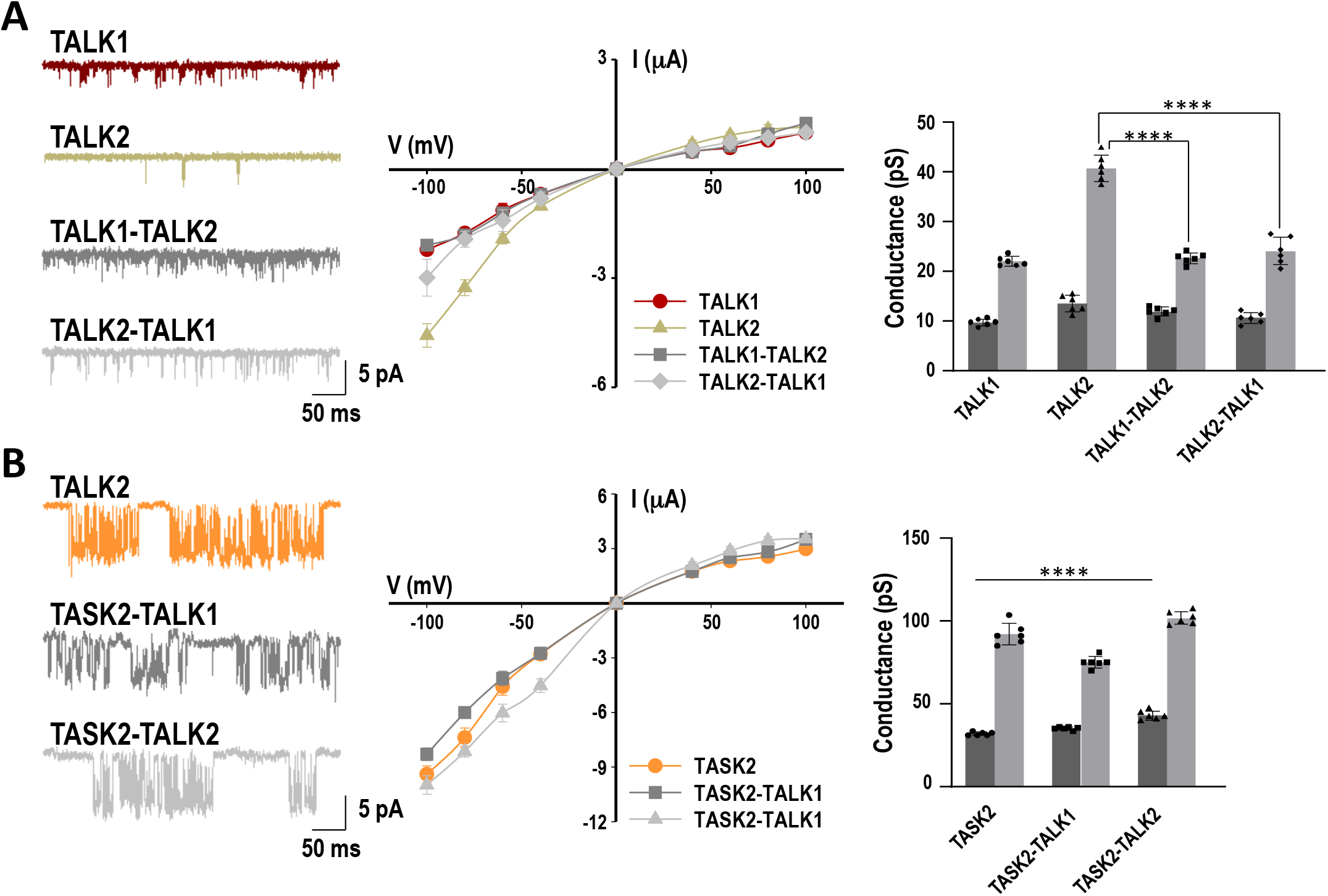
Single-channel properties of homodimeric and heterodimeric channels in HEK293 cells. (A, B) Single-channel recordings at −80 mV (left panel). Single-channel current–voltage relationships (n = 6) (middle panel). Currents were recorded in cell-attached patches held at pipette potentials from +100 mV to −100 mV in bath solution containing 150 mM KCl. Unitary conductances at −80 and +80 mV (n = 6) (Right). Data were analyzed by using unpaired t-test: ****P < 0.0001.

As observed in *Xenopus* oocytes (Fig. 4), Td-TALK1/TALK2 and Td-TALK2-TALK1 form functional channels in mammalian cells (Fig. 5A). They have unitary conductances of 12 ± 1 pS and ~11 ± 1 pS at +80 mV, and ~23 ± 1 pS and ~24 ± 3 pS at −80 mV, respectively. The tandems behave like TALK1. They have a linear current-voltage (IV) relationship whereas TALK2 shows an inward rectification, producing more currents at −80 mV than at +80 mV. Unlike TALK1 and TALK2, TASK2 opened in long bursts containing many closing events within each burst (Fig. 5B). The mean open time of TASK2 is 2.0 ± 0.3 ms at −80 mV, and the unitary conductance is 32 ± 1 pS at +80 mV and 92 ± 7 pS at −80 mV. Td-TASK2/TALK1 and Td-TASK2/TALK2 have comparable unitary conductances to each other’s and to TASK2. For all TASK2-containing channels, the current-voltage relations showed inward rectification.

These results show that heteromerization has a significant effect on single channel properties of the TALK/TASK2 channels. Association of TALK1 or TALK2 with TASK2 produces heterodimeric channels behaving more like TASK2 with higher unitary conductances and inward rectification.

## Discussion

A number of studies have focused on the heteromerization of K2P channels. These studies have produced interesting, but sometimes conflicting, results (for review see (18)). Demonstrating that heteromerization occurs under native conditions is a major challenge. Several experimental conditions are unfavorable: the low level of endogenous expression of ion channels in general, and of background K^+^ channels in particular, the absence of antibodies with affinity and selectivity allowing a specific immunolabelling of these channels, and also electrophysiological and pharmacological properties that make it difficult to distinguish between currents produced by homodimers and heterodimers. The prerequisite for heterodimer formation is the co-expression of two channel subunits not only in the same tissue but, more importantly, in the same cells of that tissue. The development of methods based on single-cell transcriptomics makes it possible to study such distribution. Among the tissues that co-express TALK1, TALK2 and TASK2, we chose the pancreas to study their expression at the single cell level. By analyzing data collected for another study, we show here that co-expression of two or three of the TALK1, TALK2 or TASK2 subunits can be detected in the same insulin-secreting β cells (Fig. 1). Their ability to form heterodimers was next studied in heterologous expression systems (Fig. 2 and 3). Two different methods were used: one based on immunolabeling of heterodimeric complexes in mammalian cells, and the second, more functional, on current poisoning using expression of a dominant negative subunit in *Xenopus* oocytes. Both methods unambiguously showed that TALK1, TALK2 and TASK2 form heterodimers. It should be noted that we did not observe the interaction between TALK2 and TASK1 that was previously reported (29). In their study, Suzuki et al. found that the TASK1 current was only partially inhibited by the coexpression of a dominant negative of TALK2. Controls were missing such as the impact of the TALK2^DN^ channel on other K_2P_ channels, or a dose-dependent inhibition of TASK1 by TALK2^DN^. They did not confirm the potential interaction by another method.

Four splice variants of TALK1 were identified, but only two of them are functional (44). The two functional variants, TALK1a and TALK1b, differ in their cytoplasmic C-ter, but their electrophysiological characteristics are indistinguishable. In the variants TALK1c and TALK1d the fourth transmembrane segment (M4) is missing. They do not induce current, and when coexpressed with TALK1a or TALK1b they do not influence the current produced by these functional variants, suggesting that M4 is essential for correct folding and/or oligomerization. Because these isoforms of TALK1 lacking M4 do not interact with TALK1a and TALK1b, they are not expected to form heterodimers with TALK2 and TASK2.

A major function of heterodimerization is to increase channel diversity by producing channels with novel electrophysiological, pharmacological or regulatory properties. Expression in *Xenopus* oocytes and mammalian cells has shown that this is the case for the TALK1, TALK2 and TASK2 heterodimers. TALK1-TALK2 channels show an intermediate sensitivity to extracellular pH, different from that of homodimeric TALK1 or TALK2 channels (Fig. 4). This suggests that each of the subunits transmits some of its properties to the heterodimer. TASK2-TALK1 and TASK2-TALK2 heterodimers have a similar sensitivity to external pH as TASK2. This is also observed with the single channels properties (Fig. 5). Both in terms of unitary conductance and rectification, TASK2-TALK1 and TASK2-TALK2 are more similar to TASK2 than to TALK1 and TALK2. This implies that the TASK2 subunit is dominant over TALK1 or TALK2 in the TASK2-TALK1 and TASK2-TALK2 heterodimers.

External pH is not the only factor that regulates TALK channels. TALK2 is also regulated by intracellular pH by a mechanism independent of that described for external pH (39,45). TALK channels have the highest level of expression in the duodenum and pancreas, where body fluids can be alkaline, suggesting that their sensitivity to alkalosis is physiologically relevant in these tissues. However, TALK2 is activated by alkaline pH outside of the range encountered physiologically. Even in pancreatic ductal cells where the apical membrane is exposed to alkaline pancreatic juice (pH 8), only a very limited amount of the TALK2 homomeric channel would be open. The combination of TALK2 with TALK1 or with TASK2 could confer to these heterodimeric channels a pH sensitivity more compatible with extracellular pH variations in the physiological range.

Beside pH, a number of other mechanisms are involved in the regulation of these channels. Binding sites for Gβγ, PIP_2_, phosphorylation and 14-3-3 were also identified in the cytoplasmic C-ter of TASK2 (46–48). On the other hand, TALK1 and TALK2 channels are regulated by NO and reactive oxygen species (ROS) such as superoxide ion (O_2_^-^) and singlet oxygen (^1^O_2_) (6). Intracellular osteopontin, a small pro-inflammatory molecule, specifically interacts with TALK1 and modulates its activity (49). Because the properties of the heterodimers cannot be inferred from the properties of the corresponding homodimers, these regulations will need to be studied on heterodimers expressed in heterologous systems. A precise knowledge of these regulations is essential to study their functional contribution in native tissues.

A number of pathological mutations and polymorphisms have been identified in TALK1 and TALK2 that generate either more active channels or channels less active with dominant negative properties (10–12). The impact of these mutations on heterodimers will also need to be studied. Finally, there is no specific pharmacology for TALK channels. Propanolol and propafenone, two beta-blockers with anti-arrhythmic effects have been shown to be responsible for a 2- to 3-fold activation of TALK2 but their effect on TALK1 and TALK2 has not been studied (50). Pyrazole derivatives, used for their analgesic properties, have also been tested on TASK2 but not on TALK channels (51). Lidocaine and bupivacaine, two local anesthetics, inhibit TASK2 (52). Although these molecules are not very specific for TASK2, they could possibly be tested on TASK2-containing heterodimeric channels provided that they are not or only slightly active on the other subunits of the heteromer (TALK1 or TALK2).

Since the ultimate goal of studying K_2P_ heteromerization is to determine the physiological roles of heterodimers, we will need to identify specific pharmacological modulators and/or to design tools capable of disrupting endogenous heterodimers, to alter their function *in vivo*. Inducible mouse models expressing dominant negative subunits such as TALK1^DN^ would be useful to study the role of homodimer and heterodimers, e.g. in endocrine and exocrine pancreatic cells and in cardiomyocytes.

## Experimental procedures

### Constructs

Human TALK1, TALK2 and TASK2 coding sequences (Ensembl accession numbers ENST00000373229.9, ENST00000373231.9, ENST00000359534.4) were inserted into pLIN a modified pGEM vector for expression in Xenopus oocytes, and pcDNA3-Zeo (Invitrogen) for expression in mammalian cells. The cDNA coding TALK1^DN^ was generated by site-directed mutagenesis using PCR and PfuTurbo DNA polymerase (Agilent). The whole cDNAs were sequenced. Tandems were constructed by overlapping PCRs and cloned into pLIN and pcDNA3-Zeo (Invitrogen). For immunocytochemistry experiments, HA (YPYDVPDYA), FLAG (DYKDDDDK) or V5 (GKPIPNPLLGLDST) tags were inserted at the C-terminus of the subunits. For *in situ* proximity ligation assay and single-channel recordings, sequences encoding TALK1, TALK2, TASK2, Td-TALK1/TALK2, Td-TALK2/TALK1, Td-TASK2/TALK1 and Td-TASK2-TALK2 were subcloned into the pIRES2-EGFP vector (Invitrogen).

### MDCK Cells Expression and Immunocytochemistry

Madin-Darby canine kidney (MDCK) cells were grown on coverslips in 24-well plates and transiently transfected with DNA plasmids using Lipofectamine 2000 (Invitrogen). At 24 h after transfection, cells were fixed and permeabilized, and channels were labeled with primary mouse anti-HA (HA-7, Sigma, 1/1000), rabbit anti-V5 (PRB-189P, Covance, 1/1000) or mouse anti-FLAG (M2, Sigma, 1/1000) antibodies, and secondary anti-mouse or anti-rabbit antibodies coupled to Alexa Fluor 488 or 594 (Invitrogen). Coverslips were mounted in Mowiol medium with DAPI on slides, and cells were imaged by confocal microscopy (LSM780 Carl Zeiss).

### Duolink Proximity Ligation Assay (PLA)

MDCK cells, grown on coverslips in 24-well plates, were transiently transfected with DNA plasmids (0.5 μg each). Cells were fixed, permeabilized, and incubated with primary mouse anti-FLAG (M2, Sigma, 1/1000) and rabbit anti-HA (Santacruz Ab, sc-805, 1/1000) or rabbit anti-V5 (PRB-189P, Covance, 1/1000) antibodies for 2 h at 37 °C in antibody diluent solution (Duolink in situ kit, Sigma-Aldrich). The cells were then labeled with the PLA antimouse PLUS probe (DUO92001) and the PLA anti-rabbit MINUS probe (DUO92005) purchased from Sigma-Aldrich. Detection was performed with Duolink in situ detection reagents red kit (DUO92008) according to the manufacturer’s protocol. Finally, the coverslips were mounted on slides, and the fields were imaged randomly using a Zeiss microscope with a 40X objective. Differences among groups were analyzed using unpaired t-test. The significance level was set at p < 0.0001. Data were represented as mean ± SEM.

### Oocyte Expression and Two-electrode Voltage Clamp Recordings (TEVC)

Capped cRNAs were synthesized using the AmpliCap-Max T7 high yield message maker kit (CellScript) from plasmids linearized by AflII. RNA concentration was quantified using a NanoDrop (Thermo scientific). Xenopus laevis stage V-VI oocytes were injected with 10-20 ng of each cRNA, and maintained at 18 °C in ND96 solution (96 mM NaCl, 2 mM KCl, 2 mM MgCl2, 1.8 mM CaCl2, 5 mM Hepes, pH 7.4). Oocytes were used 1 to 2 days after injection. Macroscopic currents were recorded with a two-electrode voltage clamp (Dagan TEV 200). The electrodes were filled with 3 M KCl and had a resistance of 0.5–2 megaohms. A small chamber with a fast perfusion system was used to change extracellular solutions and was connected to the ground with a 3 M KCl-agarose bridge. All currents were recorded in ND96. Stimulation of the preparation, data acquisition, and analysis were performed using pClamp software (Molecular Devices). All recordings were performed at 20 °C.

### Single Channel Recording

HEK293 cells were transfected with pIRES2-EGFP plasmids containing the coding sequences of TALK1, TALK2, TASK2, Td-TALK1/TALK2, Td-TALK2/TALK1, Td-TASK2/TALK1 and Td-TASK2-TALK2 using Lipofectamine 2000 and Opti-MEM medium (Life Technologies). The cells were used 2 days after transfection. Electrophysiological recording was performed using a patch clamp amplifier (Axopatch 200B, Molecular Devices). Glass patch pipettes (thick-walled borosilicate, Warner Instruments) coated with SYLGARD were used to minimize background noise. The currents were filtered at 2 kHz and transferred to a computer using the Digidata 1320 interface at a sampling rate of 20 kHz. Single-channel currents were analyzed with the pClamp program, and the plots shown in the figures were filtered at 2 kHz. For single-channel current analysis, the amplitude of each channel was set to 0.53 pA, and the minimum duration was set to 0.05 ms. In experiments using cell-attached patches, pipette and bath solutions contained (in mM): 150 KCl, 1 MgCl2, 5 EGTA, 10 glucose, and 10 HEPES (pH 7.3). All experiments were performed at 25°C. Differences among groups were analyzed using unpaired t-test. The significance level was set at p < 0.0001. Data were represented as mean ± S.D.

## Acknowledgments

We thank Martine Jodar for excellent technical assistance in molecular biology, Roberta Chiavetta and Charlotte Montillot for their help on this project during their master internship, Nathalie Leroudier for assistance in sequencing analysis, and Frederic Brau and Sophie Abelanet for support in microscopy.

## References

1. Feliciangeli, S., Chatelain, F. C., Bichet, D., and Lesage, F. (2015) The family of K2P channels: salient structural and functional properties. J Physiol 593, 2587–2603

2. Niemeyer, M. I., Cid, L. P., Gonzalez, W., and Sepulveda, F. V. (2016) Gating, Regulation, and Structure in K2P K+ Channels: In Varietate Concordia? Mol Pharmacol 90, 309–317

3. Renigunta, V., Schlichthorl, G., and Daut, J. (2015) Much more than a leak: structure and function of K(2)p-channels. Pflugers Arch 467, 867–894

4. Girard, C., Duprat, F., Terrenoire, C., Tinel, N., Fosset, M., Romey, G., Lazdunski, M., and Lesage, F. (2001) Genomic and functional characteristics of novel human pancreatic 2P domain K(+) channels. Biochem Biophys Res Commun 282, 249–256

5. Reyes, R., Duprat, F., Lesage, F., Fink, M., Salinas, M., Farman, N., and Lazdunski, M. (1998) Cloning and expression of a novel pH-sensitive two pore domain K+ channel from human kidney. J Biol Chem 273, 30863–30869

6. Duprat, F., Girard, C., Jarretou, G., and Lazdunski, M. (2005) Pancreatic two P domain K+ channels TALK-1 and TALK-2 are activated by nitric oxide and reactive oxygen species. J Physiol 562, 235–244

7. Vierra, N. C., Dadi, P. K., Jeong, I., Dickerson, M., Powell, D. R., and Jacobson, D. A. (2015) Type 2 Diabetes-Associated K+ Channel TALK-1 Modulates beta-Cell Electrical Excitability, Second-Phase Insulin Secretion, and Glucose Homeostasis. Diabetes 64, 3818–3828

8. Vierra, N. C., Dadi, P. K., Milian, S. C., Dickerson, M. T., Jordan, K. L., Gilon, P., and Jacobson, D. A. (2017) TALK-1 channels control beta cell endoplasmic reticulum Ca(2+) homeostasis. Sci Signal 10

9. Cho, Y. S., Lee, J. Y., Park, K. S., and Nho, C. W. (2012) Genetics of type 2 diabetes in East Asian populations. Curr Diab Rep 12, 686–696

10. Mahajan, R., and Gupta, K. (2014) Prevention and management of type 2 diabetes: Potential role of genomics. Int J Appl Basic Med Res 4, S1

11. Graff, S. M., Johnson, S. R., Leo, P. J., Dadi, P. K., Dickerson, M. T., Nakhe, A. Y., McInerney-Leo, A. M., Marshall, M., Zaborska, K. E., Schaub, C. M., Brown, M. A., Jacobson, D. A., and Duncan, E. L. (2021) A KCNK16 mutation causing TALK-1 gain of function is associated with maturity-onset diabetes of the young. JCI Insight 6

12. Friedrich, C., Rinne, S., Zumhagen, S., Kiper, A. K., Silbernagel, N., Netter, M. F., Stallmeyer, B., Schulze-Bahr, E., and Decher, N. (2014) Gain-of-function mutation in TASK-4 channels and severe cardiac conduction disorder. EMBO Mol Med 6, 937–951

13. Gestreau, C., Heitzmann, D., Thomas, J., Dubreuil, V., Bandulik, S., Reichold, M., Bendahhou, S., Pierson, P., Sterner, C., Peyronnet-Roux, J., Benfriha, C., Tegtmeier, I., Ehnes, H., Georgieff, M., Lesage, F., Brunet, J. F., Goridis, C., Warth, R., and Barhanin, J. (2010) Task2 potassium channels set central respiratory CO2 and O2 sensitivity. Proc Natl Acad Sci U S A 107, 2325–2330

14. Barriere, H., Belfodil, R., Rubera, I., Tauc, M., Lesage, F., Poujeol, C., Guy, N., Barhanin, J., and Poujeol, P. (2003) Role of TASK2 potassium channels regarding volume regulation in primary cultures of mouse proximal tubules. J Gen Physiol 122, 177–190

15. Warth, R., Barriere, H., Meneton, P., Bloch, M., Thomas, J., Tauc, M., Heitzmann, D., Romeo, E., Verrey, F., Mengual, R., Guy, N., Bendahhou, S., Lesage, F., Poujeol, P., and Barhanin, J. (2004) Proximal renal tubular acidosis in TASK2 K+ channel-deficient mice reveals a mechanism for stabilizing bicarbonate transport. Proc Natl Acad Sci U S A 101, 8215–8220

16. Reed, A. P., Bucci, G., Abd-Wahab, F., and Tucker, S. J. (2016) Dominant-Negative Effect of a Missense Variant in the TASK-2 (KCNK5) K+ Channel Associated with Balkan Endemic Nephropathy. PLoS One 11, e0156456

17. Toncheva, D., Mihailova-Hristova, M., Vazharova, R., Staneva, R., Karachanak, S., Dimitrov, P., Simeonov, V., Ivanov, S., Balabanski, L., Serbezov, D., Malinov, M., Stefanovic, V., Cukuranovic, R., Polenakovic, M., Jankovic-Velickovic, L., Djordjevic, V., Jevtovic-Stoimenov, T., Plaseska-Karanfilska, D., Galabov, A., Djonov, V., and Dimova, I. (2014) NGS nominated CELA1, HSPG2, and KCNK5 as candidate genes for predisposition to Balkan endemic nephropathy. Biomed Res Int 2014, 920723

18. Khoubza, L., Chatelain, F. C., Feliciangeli, S., Lesage, F., and Bichet, D. (2021) Physiological roles of heteromerization: focus on the two-pore domain potassium channels. J Physiol 599, 1041–1055

19. Czirjak, G., and Enyedi, P. (2002) Formation of functional heterodimers between the TASK-1 and TASK-3 two-pore domain potassium channel subunits. J Biol Chem 277, 5426–5432

20. Blin, S., Ben Soussia, I., Kim, E. J., Brau, F., Kang, D., Lesage, F., and Bichet, D. (2016) Mixing and matching TREK/TRAAK subunits generate heterodimeric K2P channels with unique properties. Proc Natl Acad Sci U S A

21. Blin, S., Chatelain, F. C., Feliciangeli, S., Kang, D., Lesage, F., and Bichet, D. (2014) Tandem pore domain halothane-inhibited K+ channel subunits THIK1 and THIK2 assemble and form active channels. J Biol Chem 289, 28202–28212

22. Lengyel, M., Czirjak, G., and Enyedi, P. (2016) Formation of Functional Heterodimers by TREK-1 and TREK-2 Two-pore Domain Potassium Channel Subunits. J Biol Chem 291, 13649–13661

23. Levitz, J., Royal, P., Comoglio, Y., Wdziekonski, B., Schaub, S., Clemens, D. M., Isacoff, E. Y., and Sandoz, G. (2016) Heterodimerization within the TREK channel subfamily produces a diverse family of highly regulated potassium channels. Proc Natl Acad Sci U S A 113, 4194–4199

24. Choi, J. H., Yarishkin, O., Kim, E., Bae, Y., Kim, A., Kim, S. C., Ryoo, K., Cho, C. H., Hwang, E. M., and Park, J. Y. (2018) TWIK-1/TASK-3 heterodimeric channels contribute to the neurotensin-mediated excitation of hippocampal dentate gyrus granule cells. Exp Mol Med 50, 1–13

25. Hwang, E. M., Kim, E., Yarishkin, O., Woo, D. H., Han, K. S., Park, N., Bae, Y., Woo, J., Kim, D., Park, M., Lee, C. J., and Park, J. Y. (2014) A disulphide-linked heterodimer of TWIK-1 and TREK-1 mediates passive conductance in astrocytes. Nat Commun 5, 3227

26. Lengyel, M., Czirjak, G., Jacobson, D. A., and Enyedi, P. (2020) TRESK and TREK-2 two-pore-domain potassium channel subunits form functional heterodimers in primary somatosensory neurons. J Biol Chem 295, 12408–12425

27. Plant, L. D., Zuniga, L., Araki, D., Marks, J. D., and Goldstein, S. A. (2012) SUMOylation silences heterodimeric TASK potassium channels containing K2P1 subunits in cerebellar granule neurons. Sci Signal 5, ra84

28. Royal, P., Andres-Bilbe, A., Avalos Prado, P., Verkest, C., Wdziekonski, B., Schaub, S., Baron, A., Lesage, F., Gasull, X., Levitz, J., and Sandoz, G. (2019) Migraine-Associated TRESK Mutations Increase Neuronal Excitability through Alternative Translation Initiation and Inhibition of TREK. Neuron 101, 232–245 e236

29. Suzuki, Y., Tsutsumi, K., Miyamoto, T., Yamamura, H., and Imaizumi, Y. (2017) Heterodimerization of two pore domain K+ channel TASK1 and TALK2 in living heterologous expression systems. PLoS One 12, e0186252

30. Kang, D., and Kim, D. (2004) Single-channel properties and pH sensitivity of two-pore domain K+ channels of the TALK family. Biochem Biophys Res Commun 315, 836–844

31. Decher, N., Maier, M., Dittrich, W., Gassenhuber, J., Bruggemann, A., Busch, A. E., and Steinmeyer, K. (2001) Characterization of TASK-4, a novel member of the pH-sensitive, two-pore domain potassium channel family. FEBS Lett 492, 84–89

32. Saez-Hernandez, L., Peral, B., Sanz, R., Gomez-Garre, P., Ramos, C., Ayuso, C., and Serratosa, J. M. (2003) Characterization of a 6p21 translocation breakpoint in a family with idiopathic generalized epilepsy. Epilepsy Res 56, 155–163

33. Cazals, Y., Bevengut, M., Zanella, S., Brocard, F., Barhanin, J., and Gestreau, C. (2015) KCNK5 channels mostly expressed in cochlear outer sulcus cells are indispensable for hearing. Nat Commun 6, 8780

34. Gob, E., Bittner, S., Bobak, N., Kraft, P., Gobel, K., Langhauser, F., Homola, G. A., Brede, M., Budde, T., Meuth, S. G., and Kleinschnitz, C. (2015) The two-pore domain potassium channel KCNK5 deteriorates outcome in ischemic neurodegeneration. Pflugers Arch 467, 973–987

35. Shi, Y., Stornetta, R. L., Stornetta, D. S., Onengut-Gumuscu, S., Farber, E. A., Turner, S. D., Guyenet, P. G., and Bayliss, D. A. (2017) Neuromedin B Expression Defines the Mouse Retrotrapezoid Nucleus. J Neurosci 37, 11744–11757

36. Wang, S., Benamer, N., Zanella, S., Kumar, N. N., Shi, Y., Bevengut, M., Penton, D., Guyenet, P. G., Lesage, F., Gestreau, C., Barhanin, J., and Bayliss, D. A. (2013) TASK-2 channels contribute to pH sensitivity of retrotrapezoid nucleus chemoreceptor neurons. J Neurosci 33, 16033–16044

37. Segerstolpe, A., Palasantza, A., Eliasson, P., Andersson, E. M., Andreasson, A. C., Sun, X., Picelli, S., Sabirsh, A., Clausen, M., Bjursell, M. K., Smith, D. M., Kasper, M., Ammala, C., and Sandberg, R. (2016) Single-Cell Transcriptome Profiling of Human Pancreatic Islets in Health and Type 2 Diabetes. Cell Metab 24, 593–607

38. Lesage, F., and Barhanin, J. (2011) Molecular physiology of pH-sensitive background K(2P) channels. Physiology (Bethesda) 26, 424–437

39. Li, B., Rietmeijer, R. A., and Brohawn, S. G. (2020) Structural basis for pH gating of the two-pore domain K(+) channel TASK2. Nature 586, 457–462

40. Niemeyer, M. I., Gonzalez-Nilo, F. D., Zuniga, L., Gonzalez, W., Cid, L. P., and Sepulveda, F. V. (2006) Gating of two-pore domain K+ channels by extracellular pH. Biochem Soc Trans 34, 899–902

41. Niemeyer, M. I., Gonzalez-Nilo, F. D., Zuniga, L., Gonzalez, W., Cid, L. P., and Sepulveda, F. V. (2007) Neutralization of a single arginine residue gates open a two-pore domain, alkali-activated K+ channel. Proc Natl Acad Sci U S A 104, 666–671

42. Zuniga, L., Marquez, V., Gonzalez-Nilo, F. D., Chipot, C., Cid, L. P., Sepulveda, F. V., and Niemeyer, M. I. (2011) Gating of a pH-sensitive K(2P) potassium channel by an electrostatic effect of basic sensor residues on the selectivity filter. PLoS One 6, e16141

43. Rinne, S., Kiper, A. K., Schlichthorl, G., Dittmann, S., Netter, M. F., Limberg, S. H., Silbernagel, N., Zuzarte, M., Moosdorf, R., Wulf, H., Schulze-Bahr, E., Rolfes, C., and Decher, N. (2015) TASK-1 and TASK-3 may form heterodimers in human atrial cardiomyocytes. J Mol Cell Cardiol 81, 71–80

44. Han, J., Kang, D., and Kim, D. (2003) Functional properties of four splice variants of a human pancreatic tandem-pore K+ channel, TALK-1. Am J Physiol Cell Physiol 285, C529–538

45. Niemeyer, M. I., Cid, L. P., Pena-Munzenmayer, G., and Sepulveda, F. V. (2010) Separate gating mechanisms mediate the regulation of K2P potassium channel TASK-2 by intra- and extracellular pH. J Biol Chem 285, 16467–16475

46. Anazco, C., Pena-Munzenmayer, G., Araya, C., Cid, L. P., Sepulveda, F. V., and Niemeyer, M. I. (2013) G protein modulation of K2P potassium channel TASK-2: a role of basic residues in the C terminus domain. Pflugers Arch 465, 1715–1726

47. Fernandez-Orth, J., Ehling, P., Ruck, T., Pankratz, S., Hofmann, M. S., Landgraf, P., Dieterich, D. C., Smalla, K. H., Kahne, T., Seebohm, G., Budde, T., Wiendl, H., Bittner, S., and Meuth, S. G. (2017) 14-3-3 Proteins regulate K2P 5.1 surface expression on T lymphocytes. Traffic 18, 29–43

48. Niemeyer, M. I., Cid, L. P., Paulais, M., Teulon, J., and Sepulveda, F. V. (2017) Phosphatidylinositol (4,5)-bisphosphate dynamically regulates the K2P background K(+) channel TASK-2. Sci Rep 7, 45407

49. Dickerson, M. T., Vierra, N. C., Milian, S. C., Dadi, P. K., and Jacobson, D. A. (2017) Osteopontin activates the diabetes-associated potassium channel TALK-1 in pancreatic beta-cells. PLoS One 12, e0175069

50. Staudacher, I., Illg, C., Chai, S., Deschenes, I., Seehausen, S., Gramlich, D., Muller, M. E., Wieder, T., Rahm, A. K., Mayer, C., Schweizer, P. A., Katus, H. A., and Thomas, D. (2018) Cardiovascular pharmacology of K2P17.1 (TASK-4, TALK-2) two-pore-domain K(+) channels. Naunyn Schmiedebergs Arch Pharmacol 391, 1119–1131

51. Kim, H. J., Woo, J., Nam, Y., Nam, J. H., and Kim, W. K. (2016) Differential modulation of TWIK-related K(+) channel (TREK) and TWIK-related acid-sensitive K(+) channel 2 (TASK2) activity by pyrazole compounds. Eur J Pharmacol 791, 686–695

52. Kindler, C. H., Paul, M., Zou, H., Liu, C., Winegar, B. D., Gray, A. T., and Yost, C. S. (2003) Amide local anesthetics potently inhibit the human tandem pore domain background K+ channel TASK-2 (KCNK5). J Pharmacol Exp Ther 306, 84–92

